# Combination of arsenic trioxide epigallocatechin-3-gallate and Resveratrol synergistically suppresses the growth and invasion in Brain tumor cell lines

**DOI:** 10.1101/2020.10.21.348219

**Authors:** Roshanak S. Sajjadi, Samaneh Ahmadi, Marziye Mantashloo, Sepideh Mehrpour Layeghi, Yazdan Asgari, Mohammad Ghorbani, Seyed Danial Mohammadi, Mojtaba Saffari

**Affiliations:** Cancer Biology Research Center, Cancer Institute of Iran, Tehran University of Medical Sciences, Tehran, Iran; Department of Medical Genetics, School of Medicine, Tehran University of Medical Sciences, Tehran, Iran; Department of Medical Biotechnology, School of Advanced Technologies in Medicine, Tehran University of Medical Sciences, Tehran, Iran; Department of Anatomy, School of Medicine, Tehran University of Medical Sciences, Tehran, Iran

**Keywords:** Glioblastoma, combination therapy, apoptosis, invasion, systems biology

## Abstract

The glioblastoma multiform has some properties including rapid growth, invasion, treatment resistance, and recurrence. Therefore, new therapies need to be developed that can be approved for using in patients. The previous study showed Arsenic Trioxide inhibits aggressive behavior in glioblastoma cells. Also, (–)-epigallocatechin-3-gallate prevents cellular proliferation, and invasion in multiple glioma cells. Resveratrol decreases cellular proliferation, induces cell death, and impaired the invasiveness of glioma cells. Combination therapy to inhibit cancer cells may have important clinical implications. Therefore, to assess the combination therapy of 2μM Arsenic trioxide, 100μM EGCG, and 100μM Resveratrol, we examined the metabolic activity, colony formation, media pH, cell proliferation, Caspase 3 activity, and gene expression analysis of BCL2, Caspase 3, MMP2, MMP9, CA9, u-PA, u-PAR, and Cathepsin B genes in apoptosis and invasion by both quantitative PCR experiments and Western blot assay. Systems biology tools also were used to obtain, the related network, involved pathways, and identifying the key genes in our selected criteria. The results of the study confirmed that the combined therapy prevents cell proliferation and induces of apoptosis in the Brain tumor cell lines including: U87-MG, A-172, and 1321N1. Furthermore, over-expression of caspase-3 and down-regulation of BCL-2, MMP-2, and MMP-9 confirmed that combination therapy leads to induces apoptosis and decreases invasion. Nevertheless, the lowering of pharmacological doses and improving therapeutic efficacy through combination therapy may provide advantages to treat resistance cancer cells with lower side effects. Finally, the results might suggest new modality for Glioblastoma treatment.

## 1. Introduction

Glioblastoma is the most lethal and highly aggressive brain tumor and is characterized with invasion and migration potential resulting in poor prognosis and conventional modalities, including surgery, combined radiation/chemotherapy and adjuvant chemotherapy show less promising result for the patients, with median survival 6-15 months (Ozdemir-Kaynak, Qutub et al. 2018). ATO (arsenic trioxide) that is approved by FDA, is used to treat a variety of diseases such as acute promyelocytic leukemia (APL) and successful trials with relapsed APL have contributed to an increased interest in investigating the effects of ATO in other tumor forms(Moloudi, Neshasteriz et al. 2017). However, ATO may be present in higher dose (more than 5μM) in solid tumors to be efficacious, which is higher than 2μM therapeutically dose. There has been considerable research on methods to enhance the efficacy of ATO. Drug combination therapy that has a synergistic effect, can increase efficacy of drugs(Sun, Sanderson et al. 2016). In this study, we assessed the anticancer potential of ATO combination with epigallocatechin-3-gallate which is major component of green tea and potent antioxidant of dietary phytochemicals and Resveratrol. (-)-Epigallocatechin-3-gallate (EGCG) that is the most powerful catechin found in the green tea has the one of strongest antioxidant activity and anticarcinogenic effect. Inhibition of tumorigenesis and sensitization to chemotherapy drug effects of EGCG have been studied in some of chronic diseases such as different types of cancer, cardiovascular diseases, arthritis and diabetes (Stangl, Lorenz et al. 2006, Khan and Mukhtar 2013). It is increasingly appreciated in beneficial health benefits of green tea, including chemopreventive and cancer treatment efficacy (Khan and Mukhtar 2008). Resveratrol, the major polyphenolic component of grape and red wine, has a variety of bioactivities, including antioxidant, cardioprotective, anticancer, anti-inflammation, antiaging and antimicrobial activities. It has been shown to exert anticancer property in breast cancer (Xia, Deng et al. 2010), colorectal cancer(Patel, Misra et al. 2010), U251 glioma cells (Jiang, Shang et al. 2010). In the current study we investigated the effect of combination treatment ATO, EGCG, and Resveratrol on cell proliferation, metabolic activity, colony formation ability, apoptosis induced by Caspase-3 and critical gene involved in aggressive behavior of glioblastoma cell lines. We hypothesize that combination therapy would potentiate the efficacy by increasing apoptosis rate on cell lines. Therefore, it will allow the identification of the safest and most effective treatment regimen for glioblastoma. Also, we perform pathway, GO and topological analysis to get a deep insight about mechanism which involved in our network.

## 2. Materials and Methods

### 2.1 Topological analysis

To build a Protein-Protein Interaction (PPI) network, we used the STRING plugin in Cytoscape software (v. 3.6). We set maximum additional interactors on 42 to get a 50-node network and put the confidence cut-off on 0.75 to have more certainty (Shannon, Markiel et al. 2003, Szklarczyk, Gable et al. 2019). Then, we used cytoHubba, a Cytoscape plugin, to measure a special centrality to find key proteins based on the reconstructed network (Chin, Chen et al. 2014).

### 2.2 Enrichment analysis

To perform enrichment analysis we used an R package called cluster profiler (Yu, Wang et al. 2012, Team 2015, Team 2017). Gene Ontology (GO) inquiry was conducted by clusterprofiler itself but for pathway analysis, we used KEGG over-representation analysis which essentially using KEGG database but through cluster profiler package (Kanehisa and Goto 2000). Also, charts and figures related to this subject obtained from enrichplot (another R package) which included in clusterprofiler too (Yu 2018)

### 2.3 Cell Culture

The human glioblastoma cell lines, U87-MG, A-172 and 1321N1 were purchased from the National Cell Bank of Iran (NCBI) affiliated to Pasteur Institute (Tehran, Iran). The cell lines were maintained in RPMI 1640 (Invitrogen, New Zealand) supplemented with 10% fetal bovine serum (Invitrogen), 100 units/ml penicillin and 100 μg/ml streptomycin. All cells were cultured at 37°C in a 5% CO2 in a humidified CO2 incubator. The cells were then exposed to 2μM, 100μM and 100μM concentration of ATO, EGCG, and Resveratrol (Sigma, Missouri, USA) respectively, and the combination of them. In this study, to determine whether ATO, EGCG, and Resveratrol have therapeutic effects on brain tumor cell lines, the IC50 of ATO, EGCG, and Resveratrol obtained for all the tumor cell lines and results were compared with clinical achievable of previous studies.

### 2.4 MTT Assay

The inhibitory effect of drugs on the proliferation of the cell lines were assessed by the uptake of the MTT reagent by viable cells. Cells were cultured in 96-well plates (SPL, Korea) at a density of 5,000 cell/well. The cells were cultured and treated with media containing desired concentrations of ATO, EGCG, and Resveratrol. After the treatment for 24, 48 and 72h, the cells were further incubated with 100μl MTT working solution (5mg/ml) at 37°C for 4h. The untreated cells were defined as the control group. After aspirating the solution from each well, 100μL of DMSO solution was added into each well to dissolve the crystals. The absorbance of the individual wells was detected at 570nm with a Microplate Reader ((Bio-Tek Instruments, USA).

### 2.5 Colony Formation Assay

Briefly, 5000 cells were plated in dishes and cultured at 37°C for overnight to allow full attachment and then, the cells treated with the desired concentration of ATO, EGCG, and Resveratrol for a week. On the following the cells stained with crystal violet for 20 min.And further photographed. Alpha Innotech (San Leandro, CA, USA) imaging software was used to quantify the number of cell colonies.

### 2.6 Media pH Assessment

The pH of media was measured in the presence or absence of desired concentrations of ATO, EGCG, and Resveratrol and their combinations to drop out the effects of drugs on pH. This measurement was taken before and after 72h treatment. The pH of media was measured by a laboratory pH meter.

### 2.7 Cell Proliferation Assay

Cell proliferation was measured by determining the extent of 5-Bromo-2’-deoxy-uridine (BrdU) incorporation into DNA of U87-MG cells using the BrdU cell proliferation assay ELISA kit (Roche, Germany) according to the manufacturer’s protocol. Cells (5000 cells/100μl/well) were cultured for 72 hours in flat-bottom 96-well plates treated with the desired concentrations of ATO, EGCG, and Resveratrol with a plus of BrdU for 12 hours. The cells were then fixed and DNA was denatured using 200 μl of FixDenat solution provided with the kit for 30 min and blocked with 5% BSA for 30 min at room temperature. Anti-BrdU-POD monoclonal antibody added to the cells for 90 min at room temperature. The cells were washed 3 repetitions with washing buffer, the substrate solution was added and the cells were incubated for 30min in the dark. The color reaction was activated with 1M H2SO4 for 30 min and the BrdU quantity was determined at a wavelength of 450 nm using an Absorbance Mircoplate Reader.

### 2.8 Apoptosis Assay

Apoptosis assay of U87-MG treated cells was measured using the Caspase-3 assay kit (Sigma). The Caspase-3 colorimetric assay is based on the hydrolysis of acetyl-Asp-Glu-Val-Asp p-nitroanilide (Ac-DEVD-pNA) by caspase-3, resulting in the release of the p-nitroaniline (pNA) moiety. The cells were treated with ATO, EGCG, and Resveratrol for 72h. Following centrifugation at 600g for 5 min, the cell pellets were lysed in lysis buffer provided in the kit, and the lysates were centrifuged at 14,000g for 10 min. 10μg of the supernatant was incubated with 80μl of assay buffer plus 10μl the of caspase-3 substrate acetyl-Asp-Glu-Val-Asp-p-nitroanilide (Ac-DEVD-pNA) in a 96-well plate at 37°C for 90 min. Caspase-3 activity was measured in immunosorbent assay (ELISA) reader at a wavelength of 405 nm using the detection of chromophore pNA after its cleavage by caspase-3 from the labeled caspase-3 specific substrate, DEVD-pNA.

### 2.9 Quantitative real-time RT-PCR

Total RNA was extracted from U87-MG using RNeasy Mini Kit (Qiagen) as instructed by the manufacturer after 72h treatment. The 1μg of extracting RNA was reverse transcribed into cDNA using the PrimeScript 1st strand cDNA Synthesis Kit (Clontech) according to the manufacture’s specifications. Quantitative real-time RT-PCR was performed on a Rotor-Gene Q instrument (Qiagen) using SYBR Premix Ex-Taq II (TakaraBio). 5μl SYBR Green master mix (2X), 1μl of cDNA samples, 1μl of forward/reverse primers (5pmol) and 3μl of nuclease-free water were mixed in a 0.1mL real-time PCR (Qiagen) to conduct PCR in a 10μl of reaction volume. Thermal cycling consists of an initial activation step for 30s at 95 °C, 40 cycles, including a denaturation step for 5s at 95°C and a combined annealing/extension step for 20s at 60°C. The sequences of primers are listed in Table 1. Beta-actin (ACTB) was used as a reference gene and normalize the fold change in expression of each target mRNA was calculated based on 2^-ΔΔct^ comparative expression method(Schmittgen and Livak 2008).

**Table 1.**
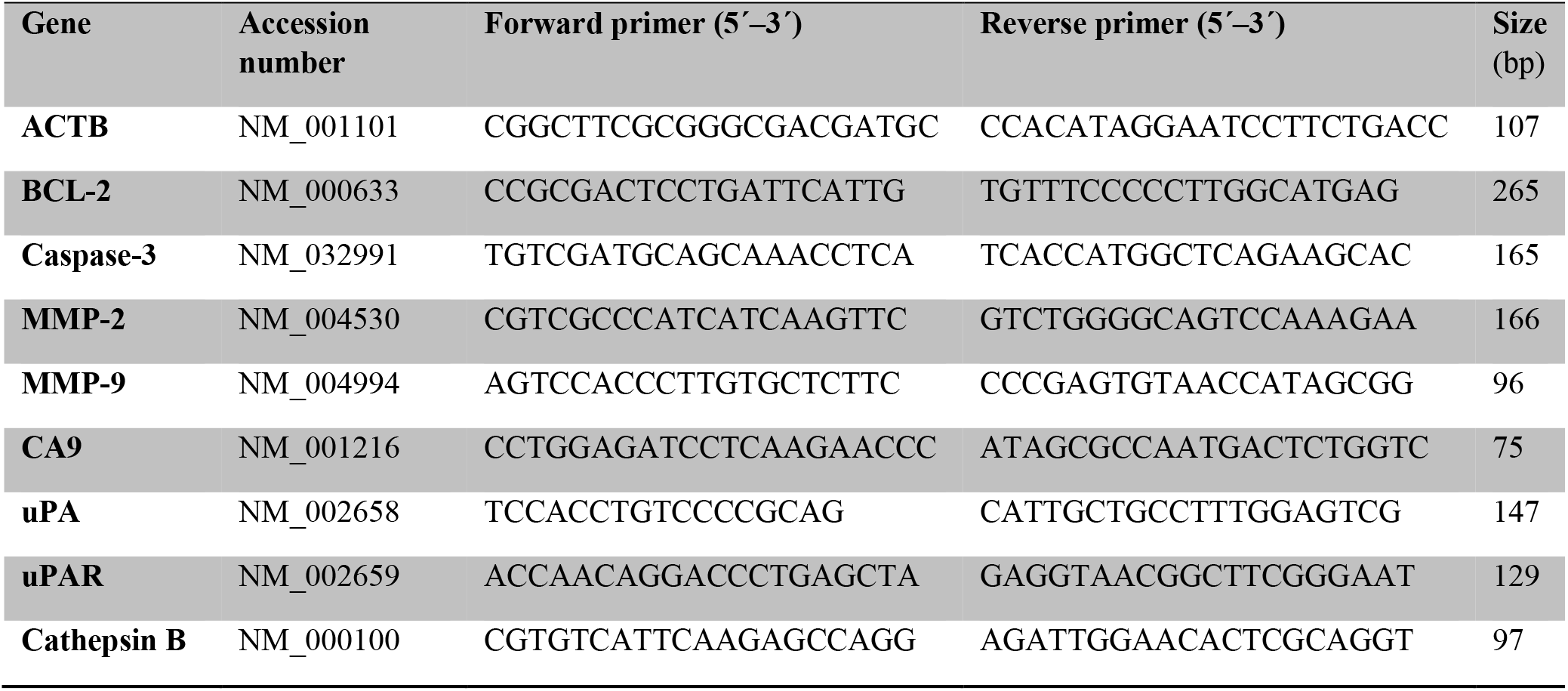
Nucleotide sequences of the primers used for real-time RT-PCR

### 2.10 Western Blot Analysis

Cells were lysed in RIPA buffer (Cell Signaling Technology, USA) supplemented with protease inhibitors cocktail (Thermo Fisher Scientific). 50μg extracted protein was separated by SDS-PAGE and transferred to PVDF membranes (BioRad Laboratories, USA). The membrane was blocked with 5% non-fat milk in TBST (10mM Tris, 150mM NaCl, 0.05%Tween 20, pH 8.3) for 1h and then probed with the indicated primary antibodies (Supplementary Table S1) at 4°C overnight with gentle shaking. The following day, the membrane was washed with TBST (5min × 3) and then incubated in m-IgGκ BP-HRP secondary antibody (Santa Cruz, sc-516102) for 1h at room temperature. Then the immunoreactive protein bands were detected using the ECL Plus reagent (Applygen Technologies, Inc., Beijing, China).

### 2.11 Data Analysis

In this study, all experiments were done in triplicate. All experimental results were processed with Microsoft Excel and Graph-Pad Prism Program 5.0 software. Data are presented as the mean ± standard deviation. The statistical analyses for the data comparisons were performed with a paired t-test. P ≤ 0.05 was considered statistically significant (*P < 0.05).

## 3. Results

### 3.1 Identifying key proteins

The reconstructed PPI network has been shown in Figure 1A which contains 50 nodes and 331 edges. To assess the importance of each component of the network, we conducted a centrality analysis. We considered two important centralities, betweenness and degree indices. The betweenness index defines as a node that is a primary connector and controls the information flow in the network. Degree index measures the number of neighbors for a node and assesses the node’s whole influence (Yu, Kim et al. 2007, Özgür, Vu et al. 2008, Tang, Kang et al. 2019). According to the results, MMP-9, MMP-2, and p53 had the best betweenness and degree scores and could be considered as the key candidate nodes through our analysis (Figure 1B).

**Figure 1.**
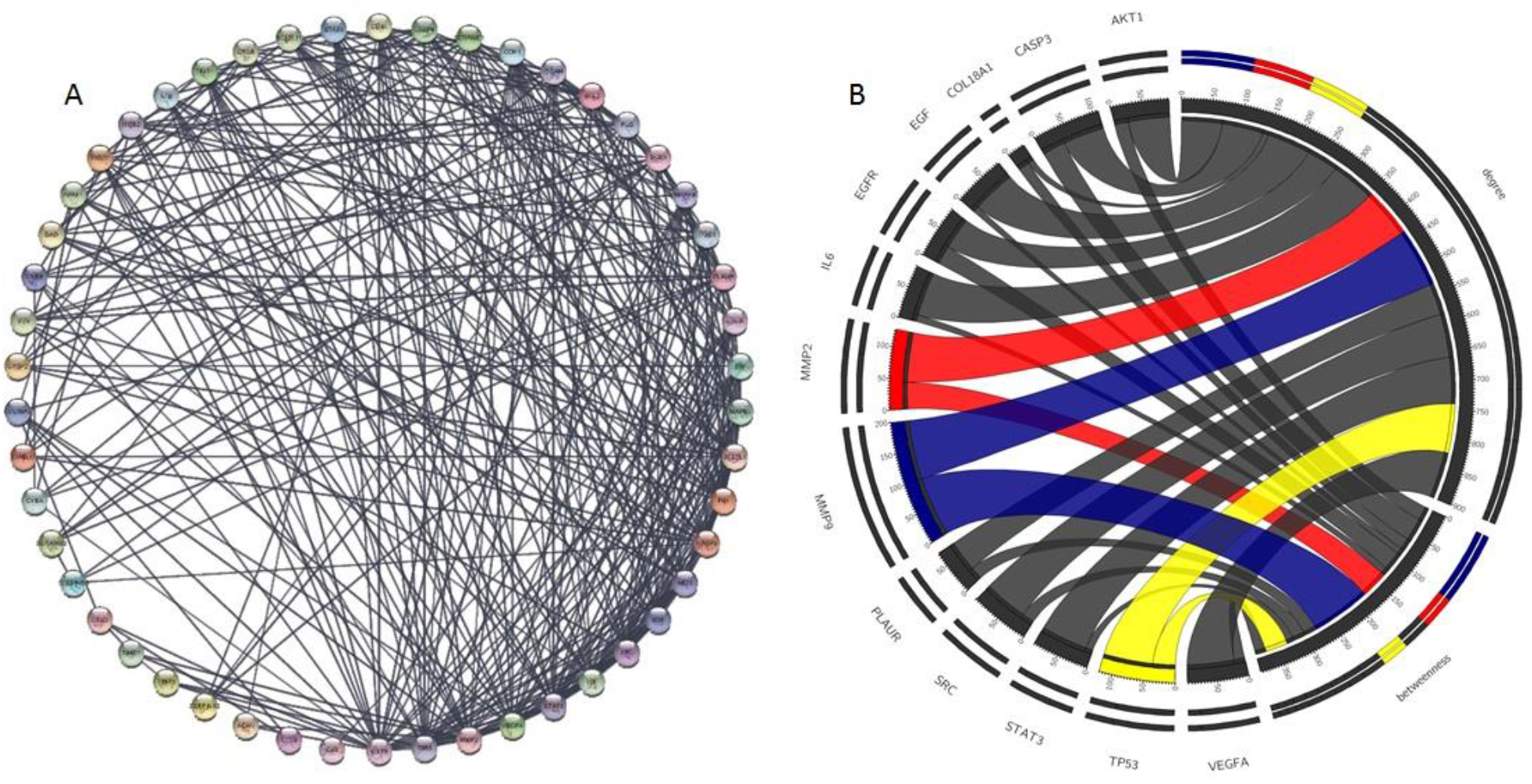
A) Reconstructed network with 50 nodes and 331 edges based on the STRING plugin. Centrality analysis identified MMP-9 and MMP-2 as proteins with the highest betweenness scores. B) Top ten proteins based on two centrality indices (betweenness and degree). As it has been shown, MMP-9, MMP-2, and TP53 are with the highest scores.

### 3.2 Enrichment analysis

According to GO biological pathway results (Figure 2A and 2B), peptidase and endopeptidase activity, apoptotic signaling and extracellular organization are in the top five results. For pathway analysis, proteoglycans in cancer and apoptosis are the most probable results (Figure 2A). One the characterization of brain cancer is irregular functions of RTKs, Proteoglycans have a significance role in this process, since they establish an interaction with extracellular ligands and their respective receptors. Proteoglycans also have a role in migration, vascularization and diffusion into the adjacent tissues in the brain (Wade, Robinson et al. 2013).

**Figure 2.**
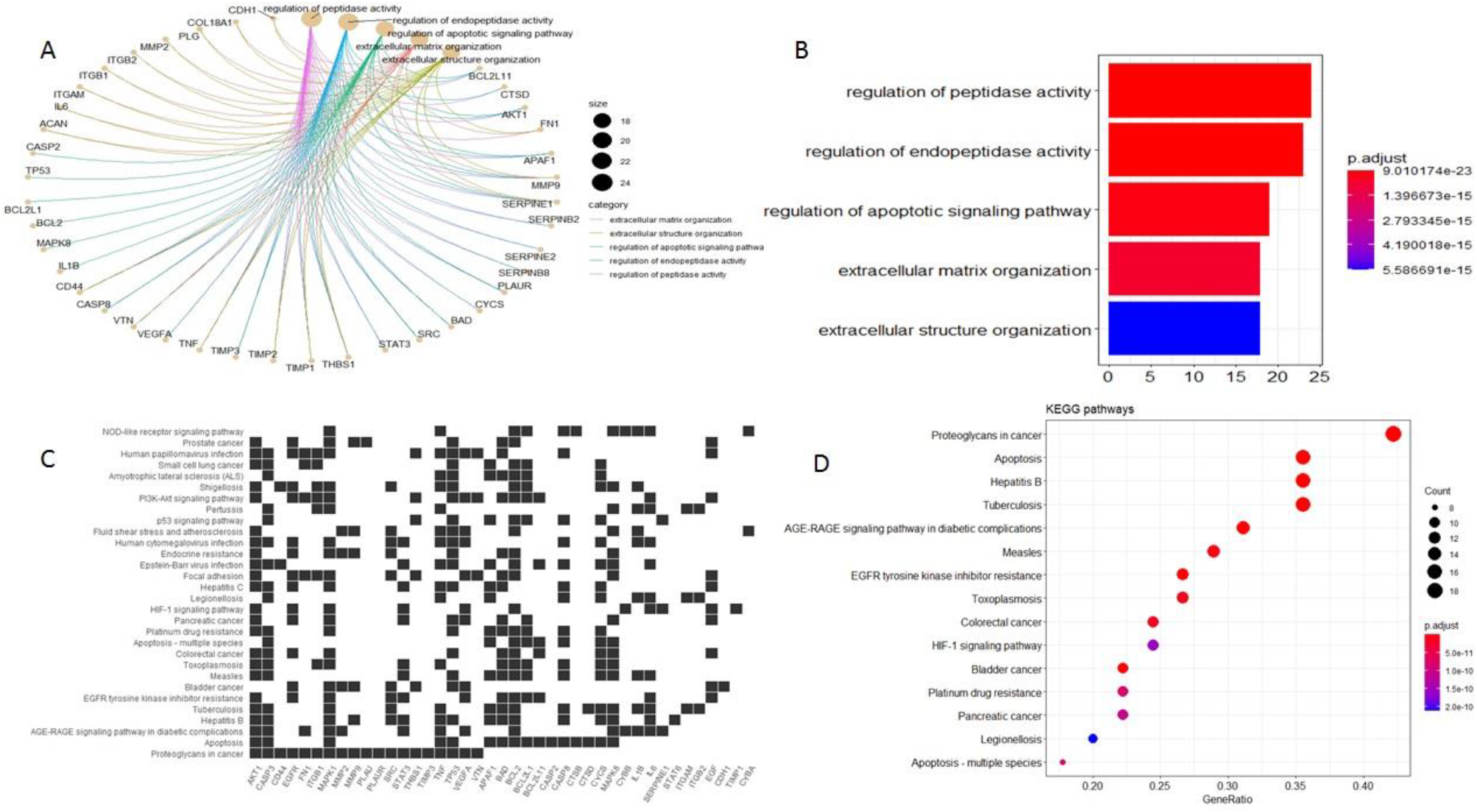
A) Cnetplot of the GO results, B) bar plot of top five of the GO terms. GO results highlighted an essential cellular process like apoptosis and extracellular matrix organization which in part accounted for migration and invasion and cell death. C) Heatmap of the pathway analysis, D) Dot plot of the KEGG analysis. Apoptosis and proteoglycans in cancer show a significant role. EGFR inhibitor and platinum drug resistance also appear in the results, which implies the significance of the reconstructed network, in which not only invasion and apoptosis processes included in the results, but also it demonstrated a drug resistance too.

### 3.3 Determination of Cellular Metabolic Activity of Cells

To determine if ATO, EGCG and Resveratrol treatment can inhibit the metabolic activity of cell lines, MTT assay was applied. The basic method is to prevent the formation of intracellular purple formazan from yellow tetrazolium compound MTT by metabolically active cells. This assay showed that the combination treatment decreased the metabolic potential of U87-MG cells more than A-172 and 1321N1 cell lines. As shown in Figure 3, the 72hrs treatment with combination of ATO (2μM), EGCG (100μM) and Resveratrol (100μM) showed significantly decreased the cell viability. After 72hrs, the combination reduced of metabolic potential by 45%, 27% and 32% on U87-MG, A-172, and 1321N1 cell lines, respectively. These reductions were not seen in treated with ATO, EGCG or Resveratrol by alone (data not shown). These results suggest that combination treatment with ATO, EGCG and Resveratrol significantly decreases cell metabolic activity.

**Figure 3.**
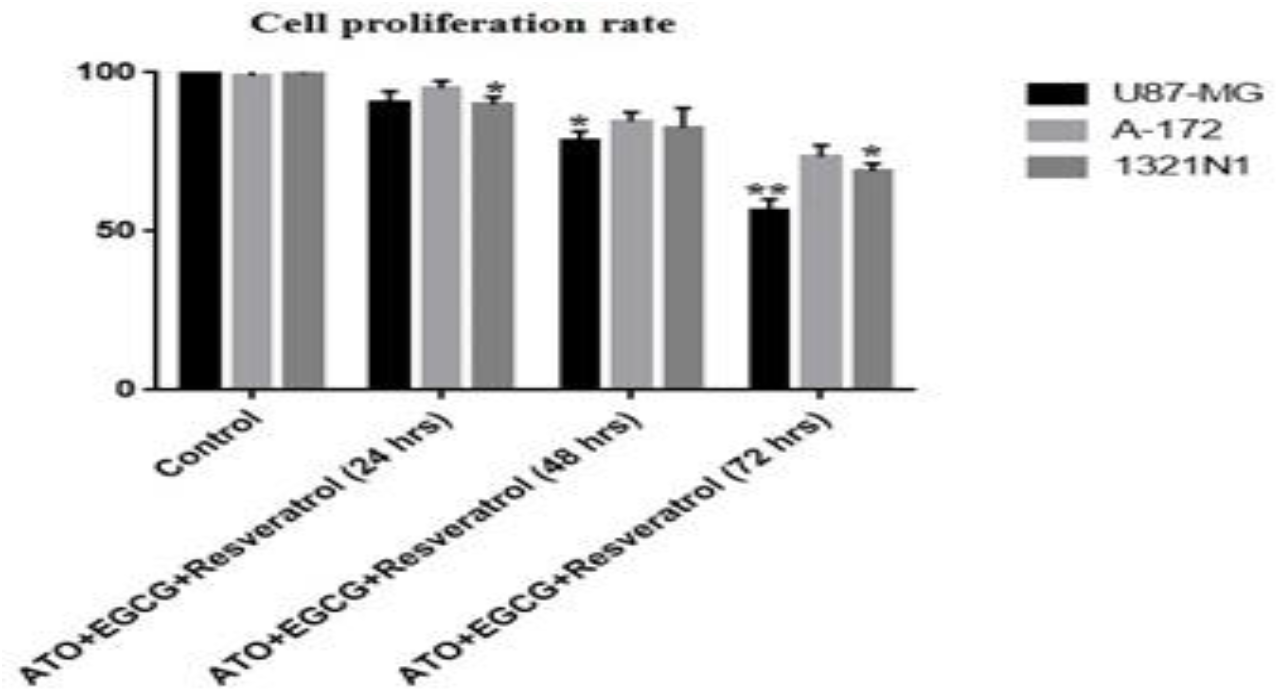
Combination treatment with ATO (2μM), EGCG (100μM) and Resveratrol (100μM) inhibits metabolic activity and cell proliferation rate in cells which treated with the mentioned concentrations. MTT assays applied in 24, 48, and 72 hours. The most reduction was seen in U87-MG cell line at 72 hours (about 45% reduction cell proliferation). The inhibitory rate was measured as percent compared to the untreated control. Values are given as mean± SD. Statistically different values of *p< 0.05 and **p<0.01 were determined compared with the control.

### 3.4 Combination Therapy Inhibits Colony Formation

On colony formation assay, showed concomitant treatment with ATO (2μM), EGCG (100μM) and Resveratrol (100μM) reduced the number of colonies to 38% on U87-MGcell line. This data suggested an enhanced antiproliferative effect of ATO, EGCG, and Resveratrol combination treatment. Table 2 shows the less and the most reduction of colony count between untreated cell line and combination treated cell line.

**Table 2.** The combined treatment with ATO, EGCG and Resveratrol decreased clonogenic survival on brain tumor cell lines.

| Group | Cell line | Number of colonies counted | Reduction of colony count compared to control (%) |
| --- | --- | --- | --- |
| Control | U87-MG | 150 | _____ |
|  | A-172 | 135 | _____ |
|  | 1321N1 | 160 | _____ |
| ATO+EGCG+Resveratro | U87-MG | 93 | 38 |
|  | A-172 | 95 | 30 |
|  | 1321N1 | 108 | 32 |

### 3.5 Combination Treatment with ATO, EGCG and Resveratrol Alkalize Media

Acidification of extracellular pH drives protease-mediated digestion and remodeling of the extracellular matrix. Treatment with ATO (2μM), EGCG (100μM) and Resveratrol (100μM) change pH of the media significantly. As displayed in Figure 4, desired concentrations of ATO, EGCG, and Resveratrol decreased significantly pH of the media on U87-MG and 1321N1 cell line to 7.17 and 7.15 respectively.

**Figure 4.**
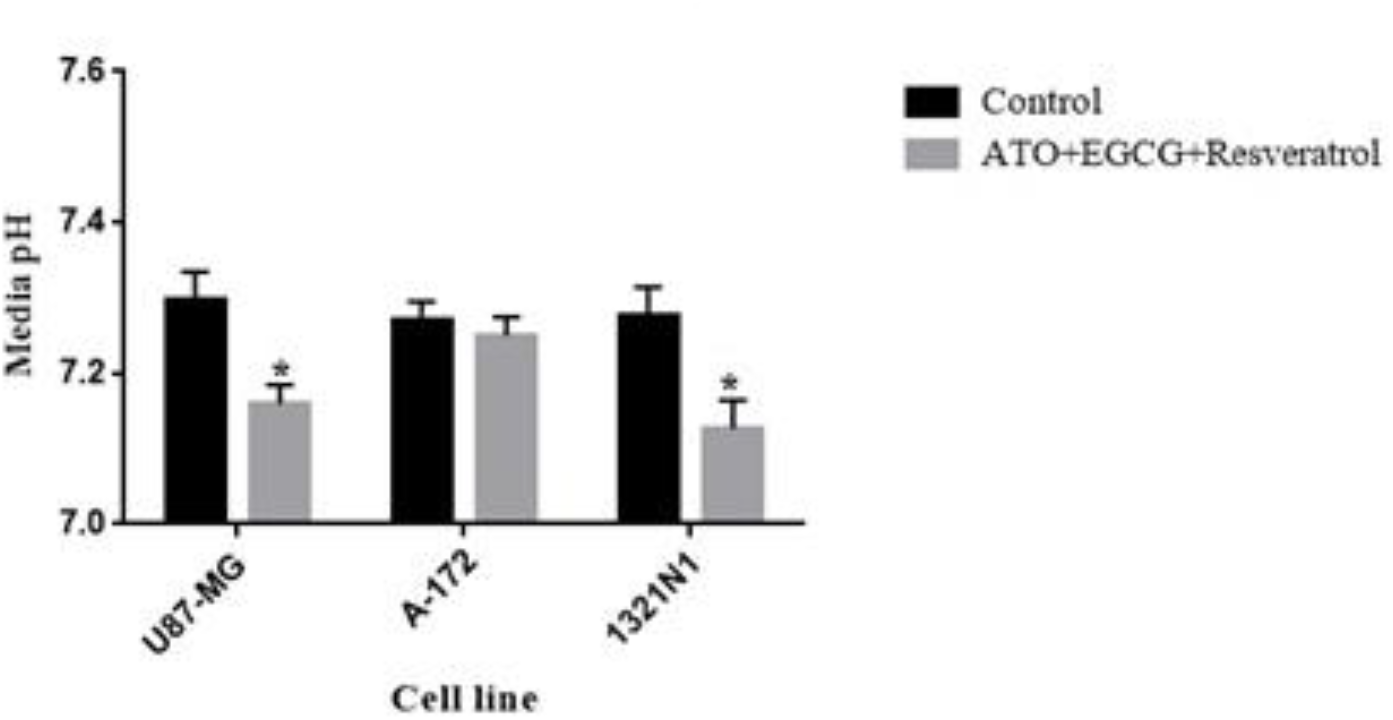
The combination treatment with ATO (2μM), EGCG (100μM) and Resveratrol (100μM) increase acidification of extracellular pH. The media pH was measured immediately after treatment with desired concentrations of ATO, EGCG and Resveratrol then was measured again after 72 hours treatment. Values are given as mean ± SD. Statistically different values of *p< 0.05 were determined compared to the control.

### 3.6 Combination Treatment Decreases Cell Proliferation

BrdU, a synthetic analogue of thymidine, is implemented into DNA during the cell cycle. The increased level of BrdU was equal to the increased cell proliferation. BrdU cell proliferation assay showed that the Resveratrol (100μM) combination with ATO (2μM), and EGCG (100μM) decreased cell proliferation by 25%, 20% and 22% on U87-MG, A-172, and 1321N1 cell lines respectively (Figure 5).

**Figure 5.**
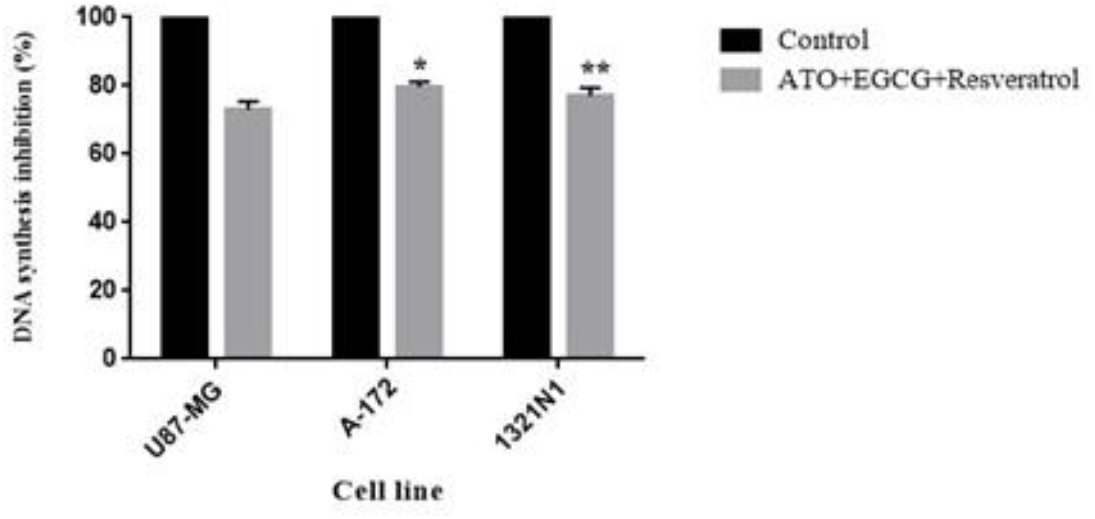
Combination treatment with ATO (2μM), EGCG (100μM) and Resveratrol (100μM) inhibits DNA synthesis rate in brain tumor cells. BrdU incorporation assays applied in 72 hours. The cell lines treated with the desired concentrations. The cell proliferation inhibition rate was measured as percent compared to the untreated control. This figure shows that the cell proliferation inhibition rate is more than 20% on U87-MG and 1321N1 cell lines. Values are given as mean± SD. Statistically different values of *p< 0.05 and **p<0.01 were determined compared with the control.

### 3.7 ATO, EGCG and Resveratrol Combination Treatment Induced Apoptotic Death

We assessed caspase-3 activity after 72 h treatment with the combination of ATO (2μM), EGCG (100μM) and Resveratrol (100μM). The results showed active caspase-3 levels were increased by17%, 15%, and 11% in U87-MG, A-172, and 1321N1 cell lines respectively. This results indicate that the combined treatment with ATO, EGCG and Resveratrol induces apoptotic cell death, probably through the activation of caspase-3 (Figure 6).

**Figure 6.**
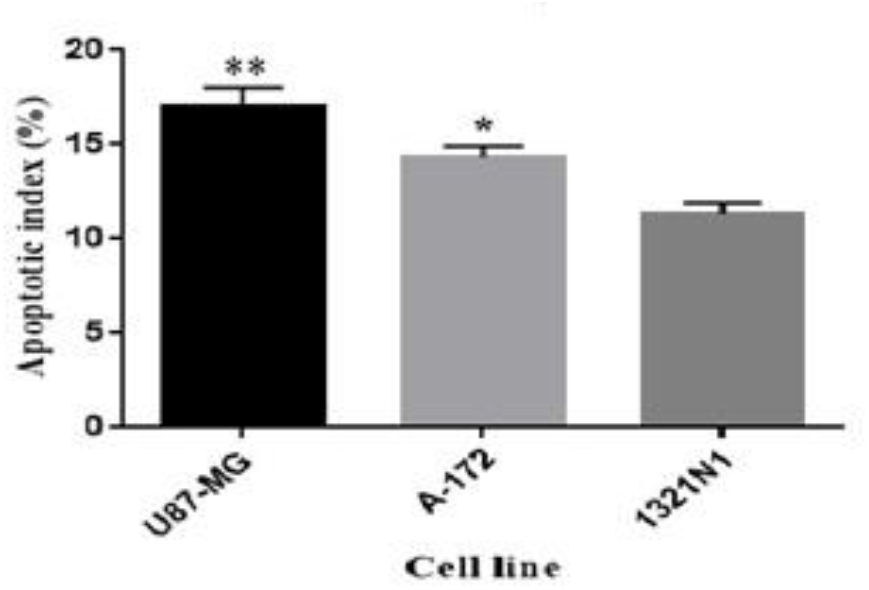
ATO (2μM), EGCG (100μM) and Resveratrol (100μM) combination induce cell death. Cells were treated for 72 hrs. The cell lysates were exposed to the peptide substrate acetyl-Asp-Glu-Val-Asp p-nitroanilide (Ac-DEVD-pNA) for 2 hours at 37°C. Fold changes in caspase-3 activity were evaluated by measuring the concentrations of p-nitroanilide (p-NA) released from the substrate due to enzymatic activity of caspase-3. Data showed that the most apoptotic rate was seen in U87-MG treated with ATO, EGCG, and Resveratrol (about 17%). Values are given as mean ± SD. Statistically different values of *p< 0.05 and **p<0.01 were determined compared to the control.

### 3.8 Combination therapy suppresses mRNA level of MMP2, 9, uPA, uPAR and Cathepsin B and induces BCL2 and Caspase 3

To measure the effect of desired combination therapy on apoptosis and invasion we investigated the relative expression level of BCL2, Caspase 3, MMP2, MMP9, CA9, u-PA, u-PAR, and Cathepsin B genes in apoptosis and invasion pathway. According to figure 7, in the combined treatment of ATO (2μM), EGCG (100μM) and Resveratrol (100μM), the least expression level of BCL2, an anti-apoptotic gene, was seen in U87-MG cell line. On the other hand, the most expression of the proapoptotic gene caspase-3 was shown just in U87-MG. Also, the significant reduction in the expression levels of MMP2, MMP9, CA9, u-PA, u-PAR and Cathepsin B genes were seen in U87-MG treated by ATO+EGCG+Resveratrol combination.

**Figure 7.**
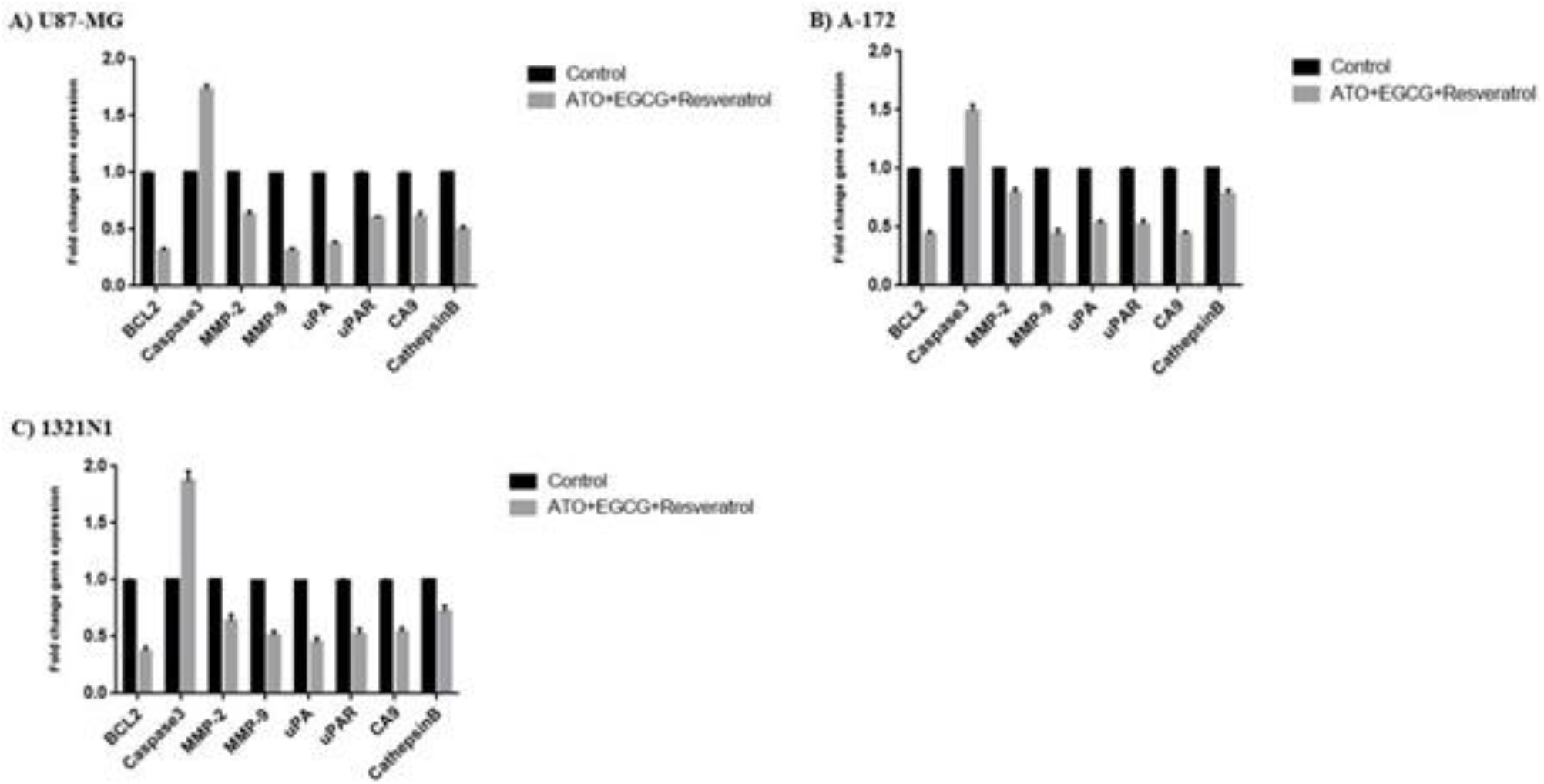
Transcriptional effect of combination treatment with ATO (2μM), EGCG (100μM) and Resveratrol (100μM) on BCL2, Caspase-3, MMP-2, MMP-9, CA-9, uPA, uPAR, and Cathepsin B genes on three brain tumor cell lines. The relative mRNA expression of each gene was measured using RT-qPCR in treated cells after normalizing the cycle thresholds of each triplicate against their corresponding ACTB. Values are given as mean± SD. Statistically different values of P< 0.05, were determined compared with the control.

### 3.9 Downregulation of MMP-2 and MMP-9 by ATO, EGCG and Resveratrol Combination Treatment

With regards to results of the expression level of desired genes, it has been determined that in U87-MG cell line, the expression levels of MMP-2 and MMP-9 had the least expression level in this three cell lines. Therefore, we investigated the protein expression level of MMPs just in U87-MG cell line. For tumor invasion, there are different types of proteases which involve in ECM degradation in tumor. Western blot analysis was used to detect the protein expression levels of MMP-2 and MMP-9 in U87-MG cell line. According to figure 8, the combined therapy with ATO (2μM), EGCG (100μM) and Resveratrol (100μM) has been decreased the protein expression levels of MMP-2 and MMP-9 after 72 hours.

**Figure 8.**
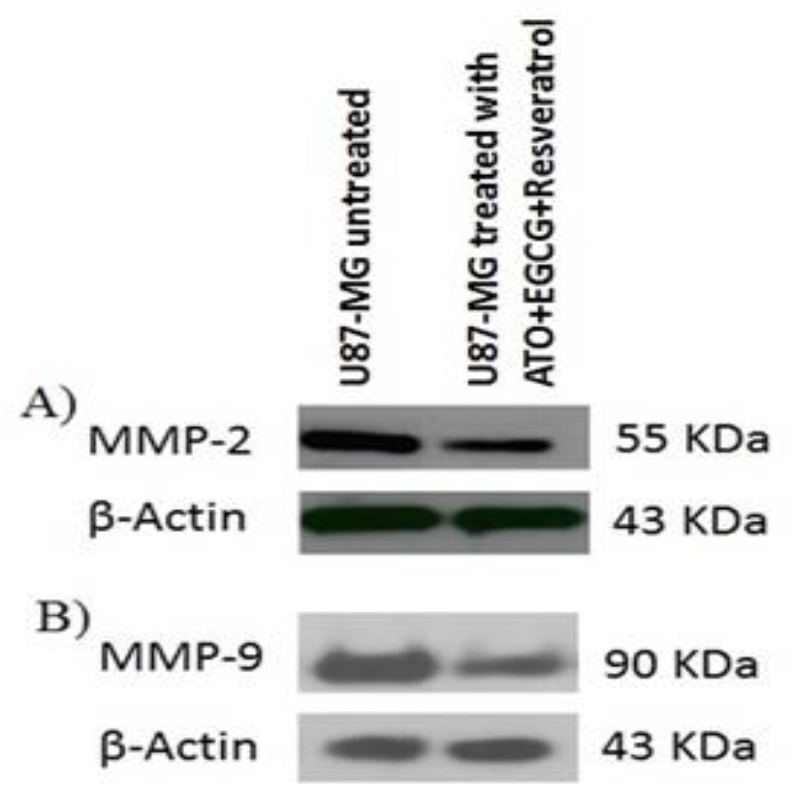
Effect of ATO, EGCG and Resveratrol combination on invasion and migration of U87-MG cell line. Western blot analysis indicated that the combined treatment with ATO (2μM), EGCG (100μM) and Resveratrol (100μM) inhibited the protein level of MMP-2 and MMP-9 compare to untreated cell line after 72 hours.

## 4. Discussion

Natural dietary agents including fruits, vegetables and spices, have been broadly studied for great potential in preventing and treating a variety of diseases including cancers and chronic diseases (Aggarwal and Shishodia 2006, Meeran, Ahmed et al. 2010). These dietary agents have little or no toxic effects on normal cells, they cost low, and they can be consumed orally. Multiple molecular targets of dietary agents have been identified (Aggarwal and Shishodia 2006, Lee, Bode et al. 2011), including the regulation of various cellular processes, such as cell proliferation, cell cycle, cell survival, and apoptosis which impact on cancer development and progression (Yoon and Baek 2005). According to the previous study, we selected doses of ATO, EGCG and Resveratrol based on plasma achievability which used to increase the effect of ATO on glioblastoma U87-MG cells (Shen, Chen et al. 1997, González-Vallinas, González-Castejón et al. 2013). Significant reduction of cell viability has been shown by the combined treatment of ATO, EGCG, and Resveratrol by measuring cell proliferation, DNA synthesis and caspase-3 activity in U87-MG, A-172, and 1321N1 cell lines. Although, the most downregulation of BCL-2 expression has been shown in U87-MG cell line treated with the combination of ATO, EGCG, and Resveratrol, but the most overexpression of caspase 3 were significantly seen in 1321N1 cell line. It has been demonstrated that BCL-2 lead to chemo-/radioresistance to glioblastoma cells and also the overexpression of BCL-2 cause to invasion and migration in human glioma cells (Wild-Bode, Weller et al. 2001, Xia, Jiang et al. 2018). Cathepsin B, a lysosomal cysteine protease, is involved in the migration and invasion of the human glioblastoma cells and correlates with the malignant progression of astrocytoma (Gondi and Rao 2013, Aggarwal and Sloane 2014). Our data showed that the mRNA level of cathepsin B was dramatically reduced by ATO and its combinations with EGCG and Resveratrol, but EGCG and Resveratrol by alone, and or their combination did not reduce cathepsin B level significantly. Attention should be paid that the most expression level of cathepsin B were seen in the most malignant cell line, U87-MG. The 1321N1 cell line showed a little bit more expression than A-172 cell line. On the other hand, the combined treatment of ATO, EGCG, and Resveratrol veratrol decrease the expression level of cathepsin B and it may inhibit migration of U87 glioblastoma cell line. It has been showed that tumor cells have an alkaline intracellular pH (pHi) and acidic interstitial extracellular pH (pHe) values. A universal characteristic of solid tumors is the acidification of extracellular pH (pHe). Driving protease-mediated digestion, disrupting cell-matrix interaction and increasing migration of cancer cells develop by the acidic tumor microenvironment which it could play an important role in promoting dormant metastasis (Raghunand and Gillies 2000, Si, Huang et al. 2014). Carbonic anhydrase IX (CA9) and Na+-H+ exchanger (NHE1) are two important keys for the cancer cell microenvironment acidification and the expression level of them has been shown in cancer cells and their increases are related to tumor invasiveness and metastasis. Another it described in several tumors including malignant gliomas, mammary sarcoma and breast cancer (Swietach, Vaughan-Jones et al. 2007, Chiche, Ilc et al. 2009). In this study we saw pHe significant increase in 1321N1, U87-MG, and A-172 treated with ATO, EGCG, and Resveratrol combination respectively. Although however, the expression level of CA9 decreased in A-172, 1321N1, and U87-MG treated with ATO, EGCG, and Resveratrol combination respectively. It has been made clear that matrix metalloproteinases (MMPs) play key role in the invasion of glioblastoma (Manini, Caponnetto et al. 2018) and it is obvious that MMP-2 and MMP-9 that are secreted by glioblastoma multiform cells correlate with the progression of gliomas (Choe, Park et al. 2002). The previous study demonstrated that EGCG and Resveratrol and ATO can inhibit MMP-2 expression in U87-MG cells (Annabi, Bouzeghrane et al. 2005, Lin, Kuo et al. 2008). Our topological analysis also demonstrated that MMP-9 and MMP-2 could be considered as hub-genes/proteins. Besides, they contained the highest betweenness score which means they have a certain level of control in the reconstructed network which mainly consists of apoptotic and invasive related pathways. In this study, our results showed that the lower concentration of ATO, EGCG, and Resveratrol combination treatment reduces MMP-2 and MMP-9 expression and their enzymatic activities and thus the combination therapy can be used as an alternative strategy for potentiation chemo-/radiotherapy effects. Urokinase-type plasminogen activator uPA (a serine protease) and its receptor uPAR have a key role in invasion and neovascularization in glioblastoma and increased activity of uPA and uPAR is associated with a poor prognosis. The uPA antisense resulted in a loss of invasive capacity and a reduction of active forms of PI3K and Akt (Gondi, Lakka et al. 2004, Hatoum, Mohammed et al. 2019). The serine protease uPA can active MMPs and increase degradation of the extracellular matrix, but EGCG inhibits uPA and prevents invasion in U87 glioblastoma cell line (Le, Leenders et al. 2018). Besides, the nuclear translocation of p65 subunit of NF-κB is inhibited by Resveratrol, therefore, Resveratrol can suppress NF-κB factor activation which cause to a downregulation of the expression of uPA/uPAR genes and reduces invasion of glioma cells (Andrade, Ramalho et al. 2018). In this study, the expression level of uPA/uPAR genes has been significantly decreased by ATO, EGCG, and Resveratrol in U87-MG. For this reason, the current study suggests that the combined treatment of ATO, EGCG and Resveratrol is more effectual than the treatment with a single compound, may be to have additive effects on their common and/or unique targets.

## 5. Conclusion

The combined treatment leads to use a lower dose of pharmacological agents and improve therapeutic efficiency; thus, it seems that provide more advantages for treating resistance cancer cells with lower side effects. Nevertheless, combination therapy of ATO with EGCG and Resveratrol mitigated the aggressive behavior of U87-MG cell line at therapeutically achievable concentration through the multifaceted mechanism, so the results might suggest new modality for Glioblastoma treatment. Further studies in animal models of glioblastoma cancer are needed to confirm and increase the understanding of combination therapy of ATO, EGCG, and Resveratrol.

## Acknowledgment

We acknowledge the department of genetics, cancer research center. This work was supported by a grant from Tehran University of Medical Sciences.

## References

Aggarwal, B. B. and S. Shishodia (2006). “Molecular targets of dietary agents for prevention and therapy of cancer.” Biochemical pharmacology 71(10): 1397–1421.

Aggarwal, N. and B. F. Sloane (2014). “Cathepsin B: multiple roles in cancer.” PROTEOMICS-Clinical Applications 8(5-6): 427–437.

Andrade, S., M. J. Ramalho, M. do Carmo Pereira and J. A. Loureiro (2018). “Resveratrol brain delivery for neurological disorders prevention and treatment.” Frontiers in pharmacology 9.

Annabi, B., M. Bouzeghrane, R. Moumdjian, A. Moghrabi and R. Béliveau (2005). “Probing the infiltrating character of brain tumors: inhibition of RhoA/ROK-mediated CD44 cell surface shedding from glioma cells by the green tea catechin EGCg.” Journal of neurochemistry 94(4): 906–916.

Chiche, J., K. Ilc, J. Laferrière, E. Trottier, F. Dayan, N. M. Mazure, M. C. Brahimi-Horn and J. Pouysségur (2009). “Hypoxia-inducible carbonic anhydrase IX and XII promote tumor cell growth by counteracting acidosis through the regulation of the intracellular pH.” Cancer research 69(1): 358–368.

Chin, C.-H., S.-H. Chen, H.-H. Wu, C.-W. Ho, M.-T. Ko and C.-Y. Lin (2014). “cytoHubba: identifying hub objects and sub-networks from complex interactome.” BMC systems biology 8(S4): S11.

Choe, G., J. K. Park, L. Jouben-Steele, T. J. Kremen, L. M. Liau, H. V. Vinters, T. F. Cloughesy and P. S. Mischel (2002). “Active matrix metalloproteinase 9 expression is associated with primary glioblastoma subtype.” Clinical Cancer Research 8(9): 2894–2901.

Gondi, C. S., S. S. Lakka, D. H. Dinh, W. C. Olivero, M. Gujrati and J. S. Rao (2004). “Downregulation of uPA, uPAR and MMP-9 using small, interfering, hairpin RNA (siRNA) inhibits glioma cell invasion, angiogenesis and tumor growth.” Neuron glia biology 1(2): 165–176.

Gondi, C. S. and J. S. Rao (2013). “Cathepsin B as a cancer target.” Expert opinion on therapeutic targets 17(3): 281–291.

González-Vallinas, M., M. González-Castejón, A. Rodríguez-Casado and A. Ramírez de Molina (2013). “Dietary phytochemicals in cancer prevention and therapy: a complementary approach with promising perspectives.” Nutrition reviews 71(9): 585–599.

Hatoum, A., R. Mohammed and O. Zakieh (2019). “The unique invasiveness of glioblastoma and possible drug targets on extracellular matrix.” Cancer Management and Research 11: 1843.

Jiang, H., X. Shang, H. Wu, G. Huang, Y. Wang, S. Al-Holou, S. C. Gautam and M. Chopp (2010). “Combination treatment with resveratrol and sulforaphane induces apoptosis in human U251 glioma cells.” Neurochemical research 35(1): 152.

Kanehisa, M. and S. Goto (2000). “KEGG: kyoto encyclopedia of genes and genomes.” Nucleic acids research 28(1): 27–30.

Khan, N. and H. Mukhtar (2008). “Multitargeted therapy of cancer by green tea polyphenols.” Cancer letters 269(2): 269–280.

Khan, N. and H. Mukhtar (2013). “Tea and health: studies in humans.” Current pharmaceutical design 19(34): 6141–6147.

Le, C. T., W. P. Leenders, R. J. Molenaar and C. J. van Noorden (2018). “Effects of the green tea polyphenol epigallocatechin-3-gallate on glioma: A critical evaluation of the literature.” Nutrition and cancer 70(3): 317-333.

Lee, K. W., A. M. Bode and Z. Dong (2011). “Molecular targets of phytochemicals for cancer prevention.” Nature Reviews Cancer 11(3): 211.

Lin, T.-H., H.-C. Kuo, F.-P. Chou and F.-J. Lu (2008). “Berberine enhances inhibition of glioma tumor cell migration and invasiveness mediated by arsenic trioxide.” BMC cancer 8(1): 58.

Manini, I., F. Caponnetto, A. Bartolini, T. Ius, L. Mariuzzi, C. Di Loreto, A. Beltrami and D. Cesselli (2018). “Role of microenvironment in glioma invasion: what we learned from in vitro models.” International journal of molecular sciences 19(1): 147.

Meeran, S. M., A. Ahmed and T. O. Tollefsbol (2010). “Epigenetic targets of bioactive dietary components for cancer prevention and therapy.” Clinical epigenetics 1(3): 101.

Moloudi, K., A. Neshasteriz, A. Hosseini, N. Eyvazzadeh, M. Shomali, S. Eynali, E. Mirzaei and A. Azarnezhad (2017). “Synergistic effects of arsenic trioxide and radiation: Triggering the intrinsic pathway of apoptosis.” Iranian biomedical journal 21(5): 330.

Ozdemir-Kaynak, E., A. A. Qutub and O. Yesil-Celiktas (2018). “Advances in glioblastoma multiforme treatment: new models for nanoparticle therapy.” Frontiers in physiology 9: 170.

Özgür, A., T. Vu, G. Erkan and D. R. Radev (2008). “Identifying gene-disease associations using centrality on a literature mined gene-interaction network.” Bioinformatics 24(13): i277–i285.

Patel, V. B., S. Misra, B. B. Patel and A. P. Majumdar (2010). “Colorectal cancer: chemopreventive role of curcumin and resveratrol.” Nutrition and cancer 62(7): 958–967.

Raghunand, N. and R. J. Gillies (2000). “pH and drug resistance in tumors.” Drug Resistance Updates 3(1): 39–47.

Schmittgen, T. D. and K. J. Livak (2008). “Analyzing real-time PCR data by the comparative C T method.” Nature protocols 3(6): 1101.

Shannon, P., A. Markiel, O. Ozier, N. S. Baliga, J. T. Wang, D. Ramage, N. Amin, B. Schwikowski and T. Ideker (2003). “Cytoscape: a software environment for integrated models of biomolecular interaction networks.” Genome research 13(11): 2498–2504.

Shen, Z.-X., G.-Q. Chen, J.-H. Ni, X.-S. Li, S.-M. Xiong, Q.-Y. Qiu, J. Zhu, W. Tang, G.-L. Sun and K.-Q. Yang (1997). “Use of arsenic trioxide (As2O3) in the treatment of acute promyelocytic leukemia (APL): II. Clinical efficacy and pharmacokinetics in relapsed patients.” Blood 89(9): 3354–3360.

Si, Z., C. Huang, X. Gao and C. Li (2014). “pH-responsive near-infrared nanoprobe imaging metastases by sensing acidic microenvironment.” RSC Advances 4(98): 55548–55555.

Stangl, V., M. Lorenz and K. Stangl (2006). “The role of tea and tea flavonoids in cardiovascular health.” Molecular nutrition & food research 50(2): 218–228.

Sun, W., P. E. Sanderson and W. Zheng (2016). “Drug combination therapy increases successful drug repositioning.” Drug discovery today 21(7): 1189–1195.

Swietach, P., R. D. Vaughan-Jones and A. L. Harris (2007). “Regulation of tumor pH and the role of carbonic anhydrase 9.” Cancer and Metastasis Reviews 26(2): 299–310.

Szklarczyk, D., A. L. Gable, D. Lyon, A. Junge, S. Wyder, J. Huerta-Cepas, M. Simonovic, N. T. Doncheva, J. H. Morris and P. Bork (2019). “STRING v11: protein-protein association networks with increased coverage, supporting functional discovery in genome-wide experimental datasets.” Nucleic acids research 47(D1): D607–D613.

Tang, Z., B. Kang, C. Li, T. Chen and Z. Zhang (2019). “GEPIA2: an enhanced web server for large-scale expression profiling and interactive analysis.” Nucleic acids research 47(W1): W556–W560.

Team, R. (2015). RStudio: integrated development for R. Boston, MA: RStudio, Inc.

Team, R. C. (2017). R: a language and environment for statistical computing [Internet]. Vienna, Austria: R Foundation for Statistical Computing; 2020.

Wade, A., A. E. Robinson, J. R. Engler, C. Petritsch, C. D. James and J. J. Phillips (2013). “Proteoglycans and their roles in brain cancer.” The FEBS journal 280(10): 2399–2417.

Wild-Bode, C., M. Weller, A. Rimner, J. Dichgans and W. Wick (2001). “Sublethal irradiation promotes migration and invasiveness of glioma cells: implications for radiotherapy of human glioblastoma.” Cancer research 61(6): 2744–2750.

Xia, E.-Q., G.-F. Deng, Y.-J. Guo and H.-B. Li (2010). “Biological activities of polyphenols from grapes.” International journal of molecular sciences 11(2): 622–646.

Xia, Y., L. Jiang and T. Zhong (2018). “The role of HIF-1α in chemo-/radioresistant tumors.” OncoTargets and therapy 11: 3003.

Yoon, J.-H. and S. J. Baek (2005). “Molecular targets of dietary polyphenols with anti-inflammatory properties.” Yonsei medical journal 46(5): 585–596.

Yu, G. (2018). “Enrichplot: Visualization of functional enrichment result.” R package version 1(2).

Yu, G., L.-G. Wang, Y. Han and Q.-Y. He (2012). “clusterProfiler: an R package for comparing biological themes among gene clusters.” Omics: a journal of integrative biology 16(5): 284–287.

Yu, H., P. M. Kim, E. Sprecher, V. Trifonov and M. Gerstein (2007). “The importance of bottlenecks in protein networks: correlation with gene essentiality and expression dynamics.” PLoS Comput Biol 3(4): e59.

