## Supplementary Table S1 for "Combination of arsenic trioxide epigallocatechin-3-gallate and Resveratrol synergistically suppresses the growth and invasion in Brain tumor cell lines"

**Supplementary Table S1. Primary antibodies for Western blot**

| Name | Company | Dilution | Information |
| --- | --- | --- | --- |
| PTEN | CELL SIGNALING TECHNOLOGY | 1:1000 | Rabbit #9552 |
| INPP4B | CELL SIGNALING TECHNOLOGY | 1:1000 | Rabbit #4039 |
| Β-Actin | CELL SIGNALING TECHNOLOGY | 1:1000 | Rabbit #4970 |
